# *ZmMTOPVIB* is essential for double-strand break formation and bipolar spindle assembly during maize meiosis

**DOI:** 10.1101/2020.07.28.225250

**Authors:** Ju-Li Jing, Ting Zhang, Yu-Hsin Kao, Tzu-Han Huang, Chung-Ju Rachel Wang, Yan He

## Abstract

The programmed formation of DNA double-strand breaks (DSBs) during early meiosis is catalyzed by SPO11, a conserved ortholog to the A subunit of an archaeal topoisomerase VI (TopoVI) which functions as a hetero-tetramer comprising two A and two B subunits. An essential role of the meiotic TopoVI B subunit (TopVIB) in DSB formation has been reported in mouse, Arabidopsis and rice. Very recently, rice *MTopVIB* was revealed to have an unexpected role in meiotic bipolar spindle assembly, highlighting multiple functions of *MTopVIB* during rice meiosis. In this work, the meiotic *TopVIB* in maize (*ZmMTOPVIB*) was characterized. The *ZmmtopVIB* mutant plants exhibited normal vegetative growth but male and female sterility. DSB formation is abolished in mutant meiocytes. Despite normal assembly of axial elements, synapsis was severely affected and homologous pairing was disrupted in mutants. Importantly, we showed that bipolar spindle assembly was also affected in *ZmmtopVIB*, resulting in triad and polyad formation. Overall, our results demonstrate that *ZmMTOPVIB* plays critical roles in DSB formation and homologous recombination. In addition, the newly-discovered function of *MTOPVIB* in bipolar spindle assembly is likely conserved across different monocots.

**One-sentence summary:** The dual roles of *MTOPVIB* in regulating meiotic DSB formation and bipolar spindle assembly are evolutionarily conserved in monocot plants.

## Introduction

Meiotic homologous recombination is a crucial step for ensuring proper chromosome segregation and generating genetic diversity in eukaryotes (Keeney and Neale, 2006). During this process, induction of programmed DNA double-strand breaks (DSBs), which is catalyzed by an evolutionary conserved topoisomerase-like transesterase protein SPO11, initiates homologous recombination (Bergerat et al., 1997; Keeney et al., 1997). Through a trans-esterification reaction, the SPO11 dimer coordinately cleaves the double strands of a DNA molecule, forming an intermediate with one SPO11 molecule covalently linked with each 5’ end of the cleaved DNA strands (Keeney and Kleckner, 1995). The SPO11 proteins are then released with the single-strand DNA (ssDNA) oligos by the Mre11/Rad50/Xrs2 (MRX) complex and Sae2/CtIP (Neale et al., 2005; Symington and Gautier, 2011). The 5’ends are then resected to expose longer 3’ ssDNA tails which are subsequently associated with two recombinases, RAD51 and DMC1, to form nucleoprotein filaments that promote homologous pairing by single-strand invasion into homologous chromosomes (Bishop et al., 1992; Shinohara et al., 1992; Cloud et al., 2012). The strand invasion results in a nascent DNA joint molecule (JM), called the displacement loop (D-loop) (Hunter and Kleckner, 2001). Consequently, JM intermediates can be processed into crossovers (COs) with reciprocal exchanges between homologous chromosomes or non-crossovers (NCOs) (Allers and Lichten, 2001; Baudat and de Massy, 2007; Grelon, 2016).

DNA topoisomerases are essential for regulating DNA topology, such as decatenating/relaxing superhelicity and untangling DNA during replication, transcription and recombination (Corbett and Berger, 2003; Graille et al., 2008). DNA topoisomerases are classified into two families based on the cleavages they make on single-strand (type I) or double-strand DNA (type II), respectively (Champoux, 2001; Wang, 2002). The type II topoisomerases are predominant and can be further categorized into two subfamilies based on their structural similarity: type IIA and type IIB (Gadelle et al., 2014). The type IIA subfamily, such as DNA gyrase and topoisomerase II enzymes, are found throughout eubacteria and eukaryotic organisms (Forterre and Gadelle, 2009). The type IIB subfamily, such as topoisomerase VI (topo VI) is ubiquitous in archaea for decatenating DNA and also found in plants, needed for successful progression of the endoreduplication cycle (Bergerat et al., 1997; Hartung et al., 2002; Forterre et al., 2007). The Topo VI enzyme functions in a heterotetramer complex comprised of two A (TopVIA) and two B (TopVIB) subunits (Corbett and Berger, 2003; Gadelle et al., 2014). About two decades ago, SPO11, which shares general sequence homology with the TopVIA, was found to play a critical role in DSB formation in all eukaryotes (Bergerat et al., 1997; Keeney et al., 1997). Interestingly, the analogs of the corresponding TopVIB and their critical role in meiotic DSB formation have only recently been identified in mouse (Robert et al., 2016), *Arabidopsis thaliana* (Arabidopsis) (Vrielynck et al., 2016) and rice (Fu et al., 2016; Xue et al., 2016).

In eukaryotes, faithful chromosome segregation during cell division is mediated by spindle, i.e. a complex protein superstructure composed of microtubules and associated proteins (Heald et al., 1996; Compton, 2000; Wittmann et al., 2001; Xue et al., 2019). After nuclear envelope breakdown in plant meiosis, randomly-polarized microtubules self-organize into bi-polar spindles during metaphase I (Vernos and Karsenti, 1995; Heald et al., 1996; Gadde and Heald, 2004). A study in *Zea mays* (maize) has revealed that microtubules form around the condensed bivalents, and then self-organized into a bipolar spindles (Chan and Cande, 1998). In maize mutants defective in chromosome synapsis and pairing, such as *dsy1, dsy2*, and *afd1* meiocytes, spindle organization at metaphase I form normally, indicating that bipolar spindle formation is independent of paired kinetochores of bivalents (Chan and Cande, 1998). Therefore, it remains elusive that what molecular pathways or regulatory factors are responsible for the process of bipolar spindle assembly in maize.

Very recently, the rice *MTopVIB* analog (*OsMTOPVIB*) was found to play a key role in bipolar spindle assembly (Xue et al., 2019). The finding raises a critical question if this newly discovered role of *OsMTOPVIB* in meiosis is widely conserved across different plant species. Here, we characterized functions of *MTopVIB* in maize, one of the best model organisms for cytogenetic study. We found that normal DSB formation was disrupted in *ZmmtopVIB* mutant meiocytes, similar to observation in the maize *spo11-1* mutant (Ku et al., 2020). A deficiency of DSBs lead to defective homologous recombination and synaptonemal complex (SC) assembly, which consequently caused univalents at diakinesis. Moreover, bipolar spindle assembly was abnormal in *ZmmtopVIB* meiocytes, resulting in missegregated meiotic univalents which were aberrantly pulled by multi-polarized spindles, yielding triads or polyads with micronuclei. Therefore, our results support the notion that *MTOPVIB* not only displays a conserved function in DSB formation but also in bipolar spindle assembly among different monocots.

## Results

### Identification of *ZmMTOPVIB*

To identify a putative *MTOPVIB* gene in maize, the full-length amino acid sequence of *Arabidopsis MTOPVIB* was used as a query to search in the maize genome database (https://maizegdb.org/) by BLASTp analysis. We identified one candidate gene (*Zm00001d014728*) with the highest similarity to *Arabidopsis MTOPVIB* (*At1G60460*). By performing rapid amplification of cDNA ends (RACE), we obtained a 1,341 bp of full-length *ZmMTOPVIB* cDNA sequence, which consists of 12 exons and 11 introns (Figure 1A). The amino acid sequences of MTOPVIB from 10 different plant species obtained from NCBI were subject to phylogenetic analysis. The result revealed that two distinct clades of MTOPVIB homologs represent genetic divergence of monocots and dicots (Figure S1). In addition, alignment of MTOPVIB protein sequences from Arabidopsis, rice and maize revealed that their MTOPVIB proteins are largely conserved, especially in three primary domains (GHKL, Small Domain and Transducer) (Figure S2). We then investigated spatio-temporal expression patterns of *ZmMTOPVIB* by quantitative RT-PCR analyses and found that it was highly expressed in the developing tassel, moderately expressed in embryo, ear and endosperm, and only weakly expressed in root, stem and leaf (Figure S3).

**Figure 1.**
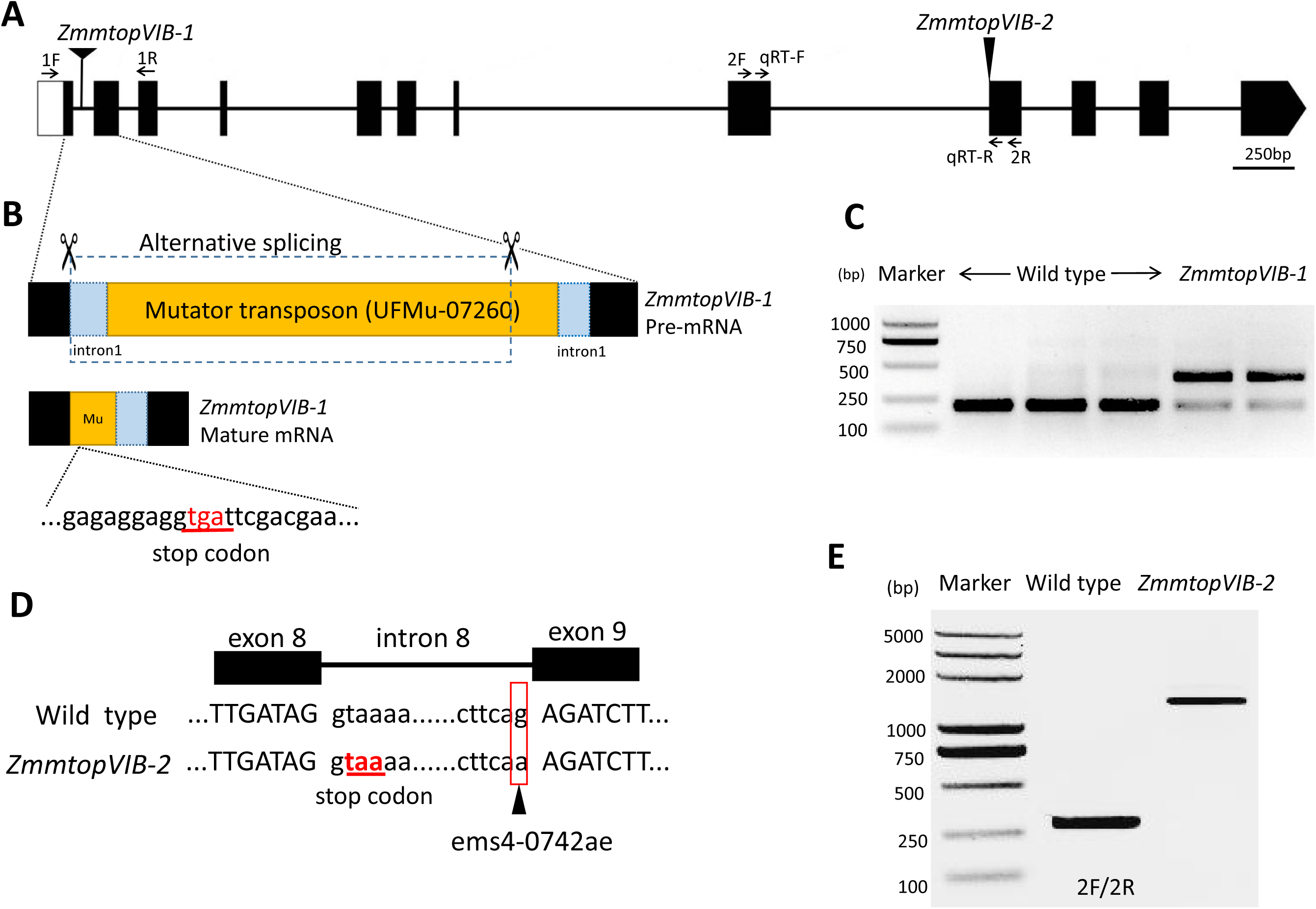
The *ZmMTOPVIB* gene and two mutant alleles. (A) Schematic diagram of maize *MTOPVIB* gene structure. Mutation sites of *ZmmtopVIB-1* and *ZmmtopVIB-2* are indicated with triangles. (B) The *ZmmtopVIB-1* mutant contains a transposon inserted in the first intron, resulting in an in-frame premature stop codon (tga). (C) RT-PCR analysis revealed two transcript variants of *ZmmtopVIB-1* using the primer pair 1F/1R. The longer transcript containing a fraction of Mutator sequence is much more abundant. (D) In the *ZmmtopVIB-2*, the EMS mutation (G to A) is located at the splicing acceptor site of the 8th intron, leading to retention of the 8th intron. The resulting transcript contains an in-frame premature stop codon (taa). (E) RT-PCR analysis confirmed the aberrant splicing transcript in the *ZmmtopVIB-2* using the primer pair 2F/2R.

To characterize functions of *MTOPVIB* in maize, we obtained one mutant line UFMu-07260 from the UniformMu population (Liu et al., 2016), which has a *Mutator* inserted in the *Zm00001d014728* gene. By PCR amplification and Sanger sequencing using the *Mutator* and *ZmMTOPVIB*-specific primers (Table S1), we confirmed that the *Mutator* transposon was inserted into the first intron of *ZmMTOPVIB* (Figure 1B). Although the insertion did not alter the *ZmMTOPVIB* expression (Figure S3), it results in aberrant splicing, leading to two major splice variants. The longer transcript appeared much abundance relative to the shorter one, which is at the same size as in the wild type (Figure 1C). Sequence analyses revealed that the longer variant contains a fraction of *Mutator* transposon sequence, resulting in an in-frame premature stop codon (underlined tga) (Figure 1B). A second mutant allele (EMS4-0742ae) was obtained from the Maize EMS-induced Mutant Database (MEMD) (Lu et al., 2018). DNA sequence analysis of the EMS4-0742ae mutant confirmed a single base mutation from G to A at the splicing acceptor site of the 8th intron (Figure 1D), which was predicted to abort splicing of the 8^th^ intron and to result in an in-frame premature stop codon (underlined taa) (Figure 1D). RT-PCR analysis showed a longer transcript in the mutant, confirming this intron retention (Figure 1E).

Both *ZmmtopVIB* mutants were completely male-sterile (Figure 2A and Figure S4A), and their anthers appeared withered at the flowering stage (Figure 2B and Figure S4B). Alexander Staining displayed that unlike the large, round, purple pollen grains of the wild type (Figure 2C and Figure S4C), mutant pollen grains were empty, shrunken, and unable to stain (Figure 2D and Figure S4D). In addition, mutant ears did not produce any seeds when pollinated with pollen from wild type plants (Figure 2E and Figure S4E). These results indicate that defective *ZmMTOPVIB* causes both male and female sterility. Hence, we named UFMu-07260 and EMS4-0742ae lines as *ZmmtopVIB-1* and *ZmmtopVIB-2* mutant alleles, respectively.

**Figure 2.**
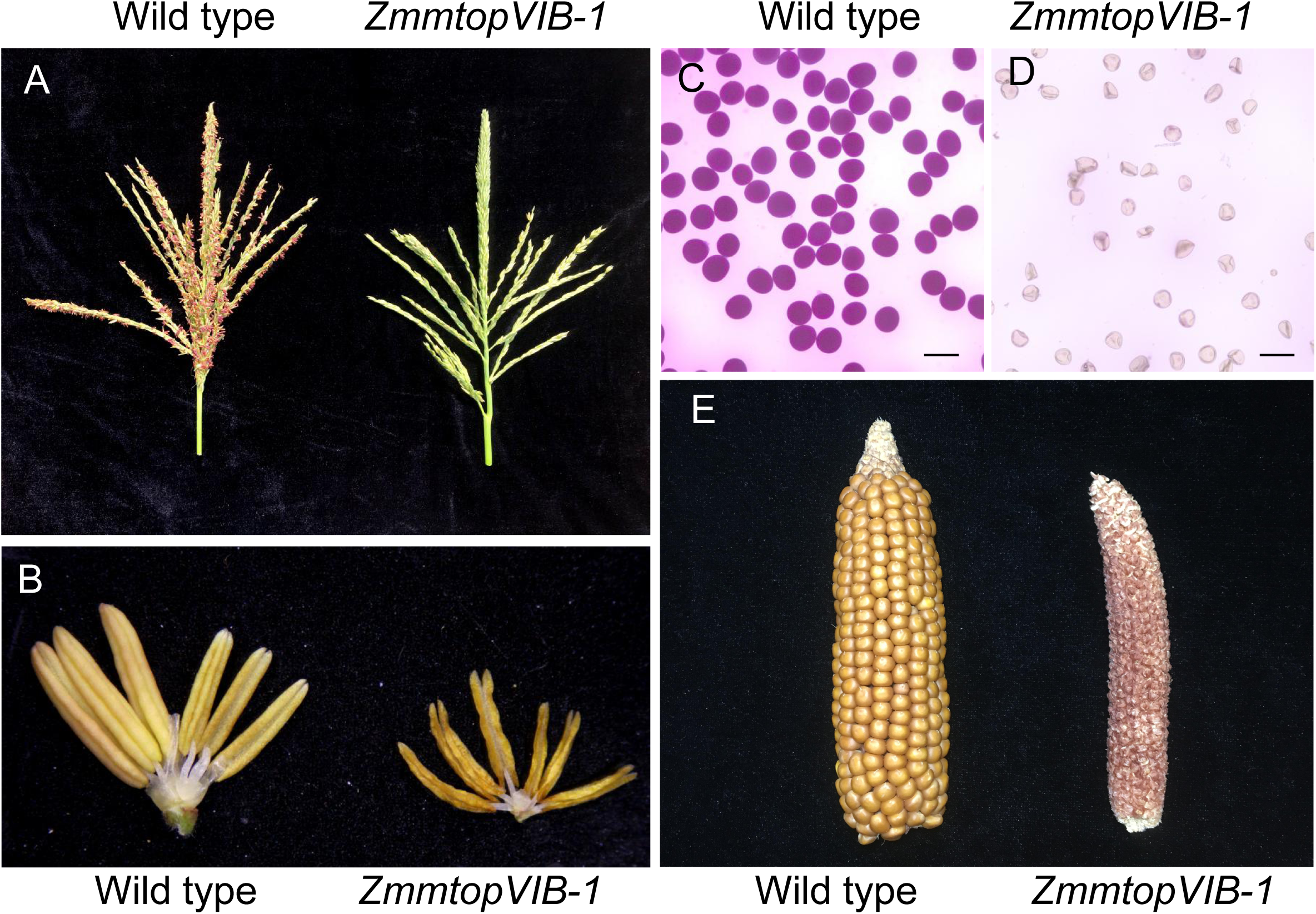
Sterile phenotypes in *ZmmtoVIB-1* mutant. (A) Tassels of wild type and mutant *ZmmtopVIB-1* at flowering stage. (B) Anthers from glume removed-spikelets of wild type and *ZmmtopVIB-1* mutant at flowering stage. (C) Mature pollen grains from wild type stained with Alexander solution. Scale bar = 100μm. (D) Sterile pollen grains from *ZmmtopVIB-1* stained with Alexander solution. (E) Harvested ears of wild type and *ZmmtopVIB-1* mutant pollinated with wild type pollen grains.

### Meiosis is disturbed in the *ZmmtopVIB* mutants

Meiotic chromosome behaviors in *ZmmtopVIB* mutants were analyzed by DAPI staining. In the wild type, long, thin, thread-like chromosomes were first observed at leptotene (Figure 3A). Then, chromosomes rearrange next to a large and off-set nucleolus and begin to pair and synapse at zygotene (Figure 3B). During pachytene, chromosomes are completely synapsed to form thick chromosome threads (Figure 3C). From diplotene to diakinesis, chromosomes condense into 10 short, rod-like bivalents that are distributed in nucleus (Figure 3D). At metaphase I, the 10 bivalents properly align on the equatorial plate (Figure 3E), and equal numbers of chromosomes move to the two opposite poles of the cell at anaphase I (Figure 3F).

**Figure 3.**
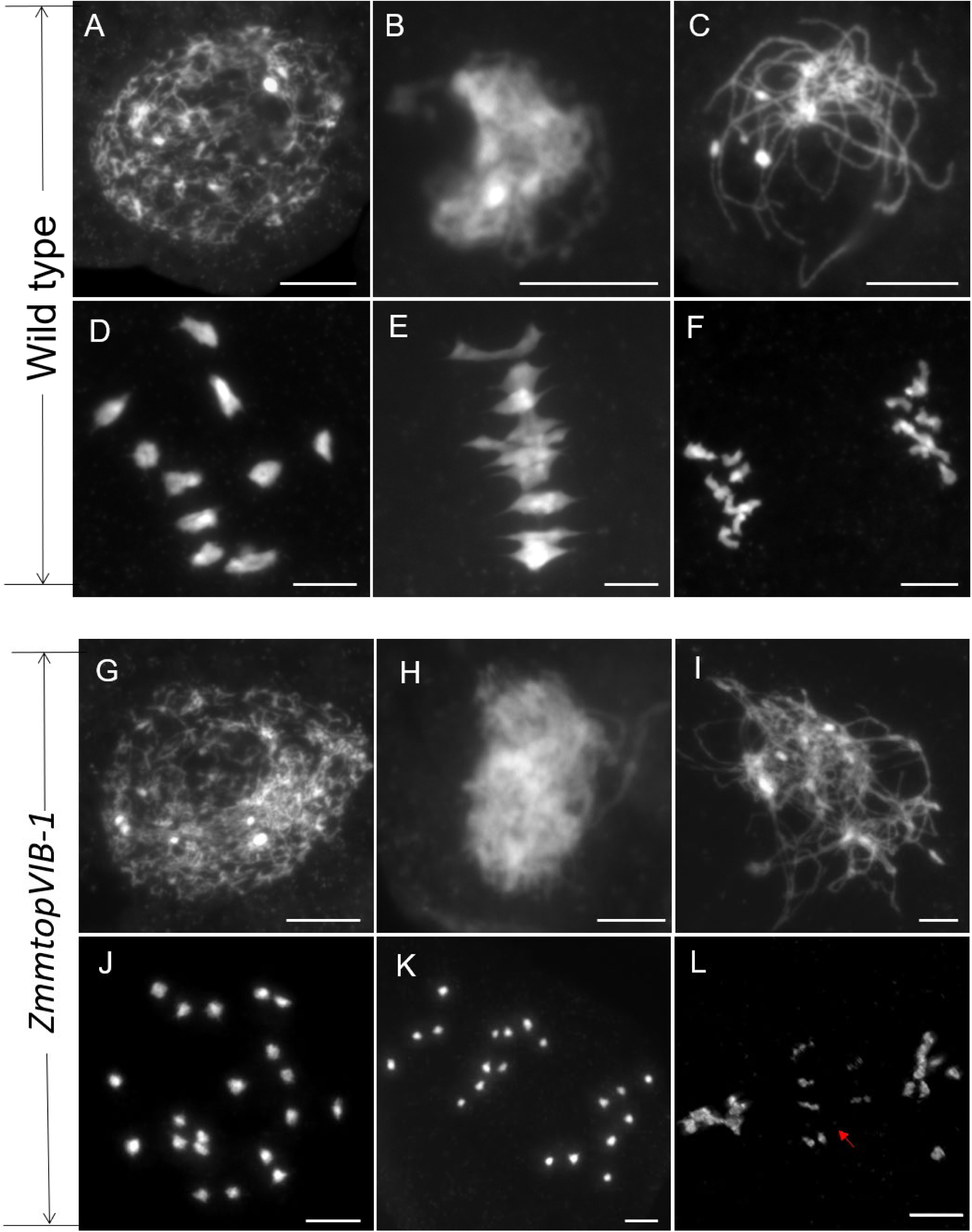
Meiotic chromosome behaviors in wild type and *ZmmtopVIB-1* mutant meiocytes. (A–F) Meiosis in wild type; (G-L) Meiosis in *ZmmtopVIB-1*. (A, G) Leptotene; (B, H) Zygotene; (C, I) Pachytene; (D, J) Diakinesis; (E, K) Metaphase I; (F, L) Anaphase I. The red arrows indicate the lagging univalents. Scale bars = 10 µm.

Chromosomal behaviors of *ZmmtopVIB-1* and *ZmmtopVIB-2* meiocytes were similar to wild type until pachytene where complete synapsis seen as thick chromosomes was not observed in mutants (Figure 3G-3I and Figure S5A-S5C). However, we observed 20 unpaired univalents randomly scattered in the nuclei of mutant meiocytes at diakinesis (Figure 3J and Figure S5D), which never aligned on the equatorial plate at metaphase I (Figure 3K and Figure S5E). At anaphase I, we recorded 20 univalents segregated randomly and asymmetrically, with lagging chromosomes often found in the center of the mutant nuclei (Figure 3L, Figure S5F). Based on these obvious abnormalities in meiotic chromosome behavior, we concluded that the sterility of the *ZmmtopVIB* mutants was due to defective meiosis. Since both *ZmmtopVIB-1* and *ZmmtopVIB-2* exhibited the similar meiotic phenotypes, all of our subsequent analyses were performed on *ZmmtopVIB-1* as a representative mutant allele.

### *ZmMTOPVIB* is essential for normal meiotic DSB formation

Formation and repair of programmed DSBs during meiosis prophase I is an essential prerequisite for homologous recombination (de Massy, 2013). Despite important roles during meiosis, DSBs represent one of most deleterious lesions, and such DNA damage rapidly induces the phosphorylation of the histone variant H2AX at its S139 residue (γH2AX) (Nakamura et al., 2010; Yuan et al., 2010). Consequently, the occurrence of γH2AX foci is routinely used as a biomarker to monitor DSB formation (Dickey et al., 2009; Lobrich et al., 2010). To investigate if DSB formation is defective in maize *ZmmtopVIB* mutants, we used an antibody that specifically recognizes γH2AX signal for immunofluorescence analysis in wild type and *ZmmtopVIB-1* meiocytes. We found that γH2AX foci appeared as punctate-like signals scattered throughout the nuclei of wild type meiocytes, reaching a peak at early zygotene and decreasing at late zygotene (Figure 4A-4C). In contrast, no γH2AX signal was detected in early zygotene of *ZmmtopVIB-1* meiocytes (Figure 4H, n=33), suggesting that normal DSB formation is defective in the *ZmmtopVIB-1* mutant. Interestingly, at late zygotene, we found a few clusters of γH2AX signals in *ZmmtopVIB-1* meiocytes, that are often associated with chromosome tangles (Figure 4I, n=32).

**Figure 4.**
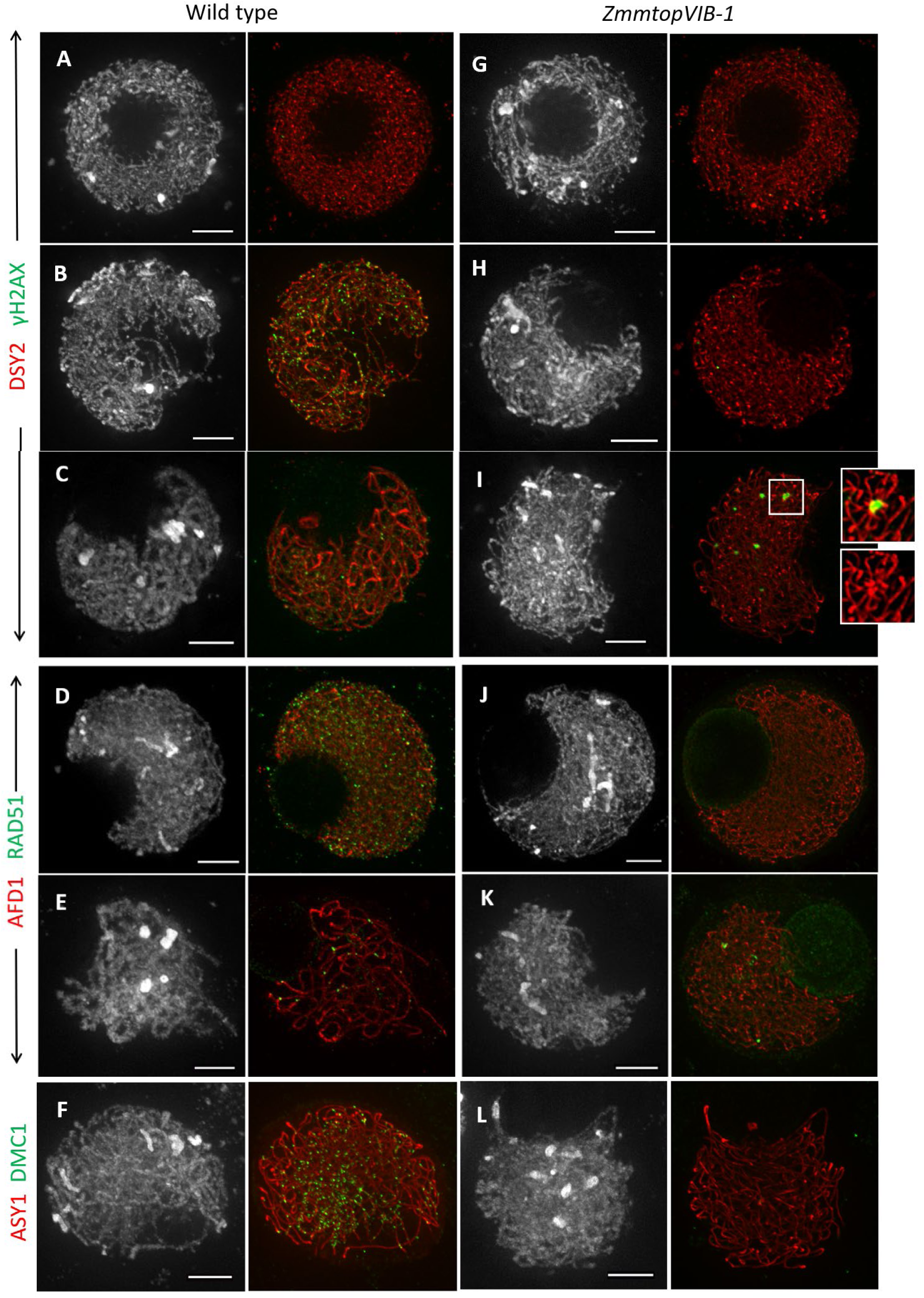
Immunolocalization of γH2AX, RAD51 and DMC1 in wild type and *ZmmtopVIB-1* meiocytes. (A-F) Wild type meiocytes with γH2AX and DSY2 signals (A-C), RAD51 and AFD1 (D-E), DMC1 and ASY1 (F). (G-L) *ZmmtopVIB-1* meiocytes with γH2AX and DSY2 signals (G-I), RAD51 and AFD1 (J-K), DMC1 and ASY1 (L). Meiocytes at leptotene (A, G), early zygotene (B, D, F, H, J, L) and late zygotene (C, E, I, L) are determined by morphology of heterochromatic knobs and lengths of anthers used for staining. Scale bars = 5 µm.

To further investigate the ectopic detection of γH2AX in mutants, we analyzed distribution of recombinases RAD51 and DMC1. While RAD51 is required for DNA repair in somatic and meiotic recombination, DMC1 functions specifically during meiotic recombination (Bishop et al., 1992; Cloud et al., 2012). In wild type meiocytes, both RAD51 (Figure 4D, n=36) and DMC1 (Figure 4F, n=52) manifested as numerous punctate foci distributed on chromosomes at zygotene. At late zygotene, RAD51 signals significantly reduced in wild type (Figure 4E). Similar to γH2AX signals, RAD51 staining was absent in early zygotene (Figure 4J, n=12), but appeared in a few foci at late zygotene (Figure 4K, n=32). Conversely, among 43 mutant meiocytes during zygotene, no obvious DMC1 signals were detected (Figure 4L). Although γH2AX signals were detected at a very low level in mutant meiocytes, they were likely DNA damages associated with chromosomal entanglements. Given the observation of absence of DMC1 signals, these aberrant DSBs may not repaired through canonical meiotic recombination pathway. Taken together, these results demonstrate that *ZmMTOPVIB* is essential for normal meiotic DSB formation in maize.

### *ZmMTOPVIB* is critical for homologous pairing but not required for telomere bouquet formation

To investigate whether defective *ZmMTOPVIB* disrupts homologous chromosome pairing, we performed fluorescence *in situ* hybridization (FISH) using the 5S rDNA probe in wild type and the *ZmmtopVIB-1* mutant. The 5S rDNA is a tandem repeat sequence located on the long arm of chromosome 2 that is routinely used to evaluate chromosome pairing and segregation in maize (Wang et al., 2018). In wild type meiocytes, two 5S signals gradually pair with each other during zygotene (Figure 5A-5B). At pachytene, the paired 5S signal was observed in all cells examined (Figure 5C, n=48). In contrast, two separate 5S rDNA signals were consistently detected in *ZmmtopVIB-1* meiocytes (Figure 5F-5G), indicating that homologous chromosome pairing is defective in the *ZmmtopVIB-1* mutant.

**Figure 5.**
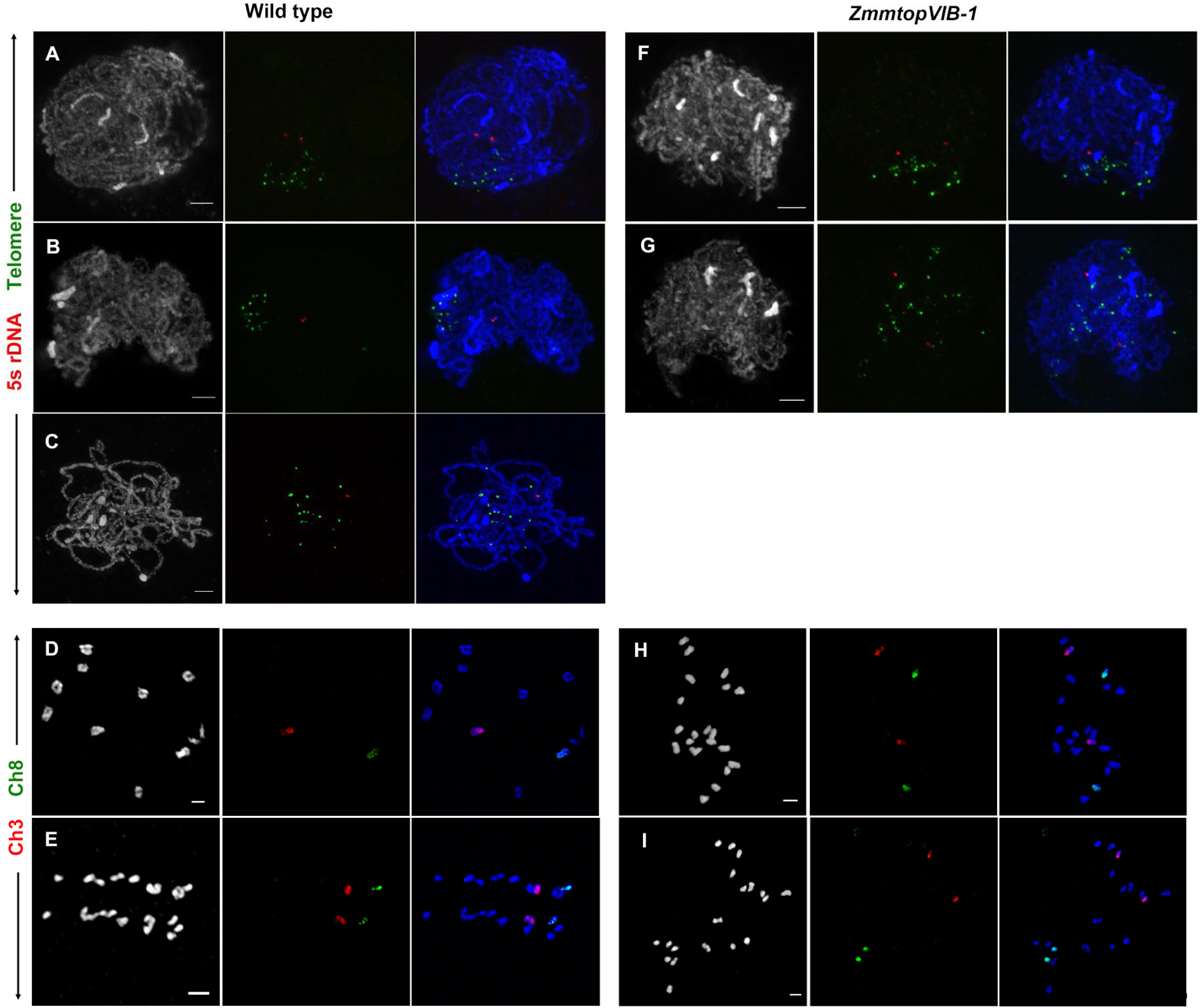
*ZmMTOVIB* is essential for homologous pairing, but not required for telomere bouquet clustering. (A-C) FISH in wild type meiocytes using a 5s rDNA (red) and telomere (green) probes at different stages. (D, E) Chromosome painting in wild type meiocytes using chromosome 3 (red) and chromosome 8 (green)-specific probes. (F, G) FISH in *ZmmtopVIB-1* meiocytes using a 5s rDNA (red) and telomere (green) probes at different stages. (H, I) Chromosome painting in *ZmmtopVIB-1* meiocytes using chromosome 3 (red) and chromosome 8 (green)-specific probes. Scale bars = 5 µm.

The telomere bouquet is an evolutionarily conserved chromosome arrangement that clusters telomeres in a small region on the nuclear envelope (Tomita and Cooper, 2007; Moiseeva et al., 2017). This specialized structure is thought to promote initiation of homologous pairing during early prophase I (Loidl, 1990; Scherthan, 2001; Harper et al., 2004). To explore whether defective *ZmmtopVIB* affects telomere bouquet formation, we conducted FISH analysis using pAtT4 probe (Richards and Ausubel, 1988; Prieto et al., 2004) in wild type and *ZmmtopVIB-1* meiocytes. In wild type (Figure 5A-5B, n=44) and *ZmmtopVIB-1* (Figure 5F, n=51) meiocytes at early zygotene stage, telomere signals were clustered and attached to the nuclear envelope, indicating that *ZmMTOPVIB* is not required for telomere bouquet formation. As mutant meiocytes did not show the normal pachytene stage, telomere signals become diffused at late zygotene (Figure 5G).

It is unexpected to observe γH2AX signals in *ZmmtopVIB* mutants. Although homologous pairing assessed by 5S rDNA signals suggested that pairing is defective, we sought to analyze more loci in the genome. Therefore, we adapted the recently developed chromosome painting method in meiosis (Albert et al., 2019). By using chromosome 3 and chromosome 8-specific probes, wild type meiocytes clearly showed ten bivalents (Figure 5D). At diakinesis, bivalents labeled with chromosome specific probe were correctly separated at anaphase I (Figure 5E). In contract, *ZmmtopVIB* mutants showed a complete failure of bivalent formation by chromosome painting (Figure 5H-5I).

### Defective installation of central element ZYP1 in *ZmmtopVIB-1* meiocytes

The synaptonemal complex (SC) is a meiosis-specific chromosomal structure that connects homologous chromosomes by a transverse filament to promote efficient CO formation (Page and Hawley, 2004; Argunhan et al., 2017). To investigate the pattern of SC localization in wild type and *ZmmtopVIB-1* mutant meiocytes, we performed immunostaining using AFD1, ASY1 and ZYP1 antibodies. Maize AFD1 is homologous to *Arabidopsis* and rice REC8 (Cai et al., 2003; Shao et al., 2011). It is a vital component of the cohesion complex associated with axial and lateral elements and is required for axial element (AE) elongation (Golubovskaya et al., 2006). AFD1 signals appeared as threads along entire chromosomes of wild type meiocytes at zygotene (Figure 6A, n=21). AFD1 signals in *ZmmtopVIB-1* meiocytes was consistent with that of wild type (Figure 6B, n=24), indicating that *ZmMTOPVIB* is not required for cohesion complex assembly.

**Figure 6.**
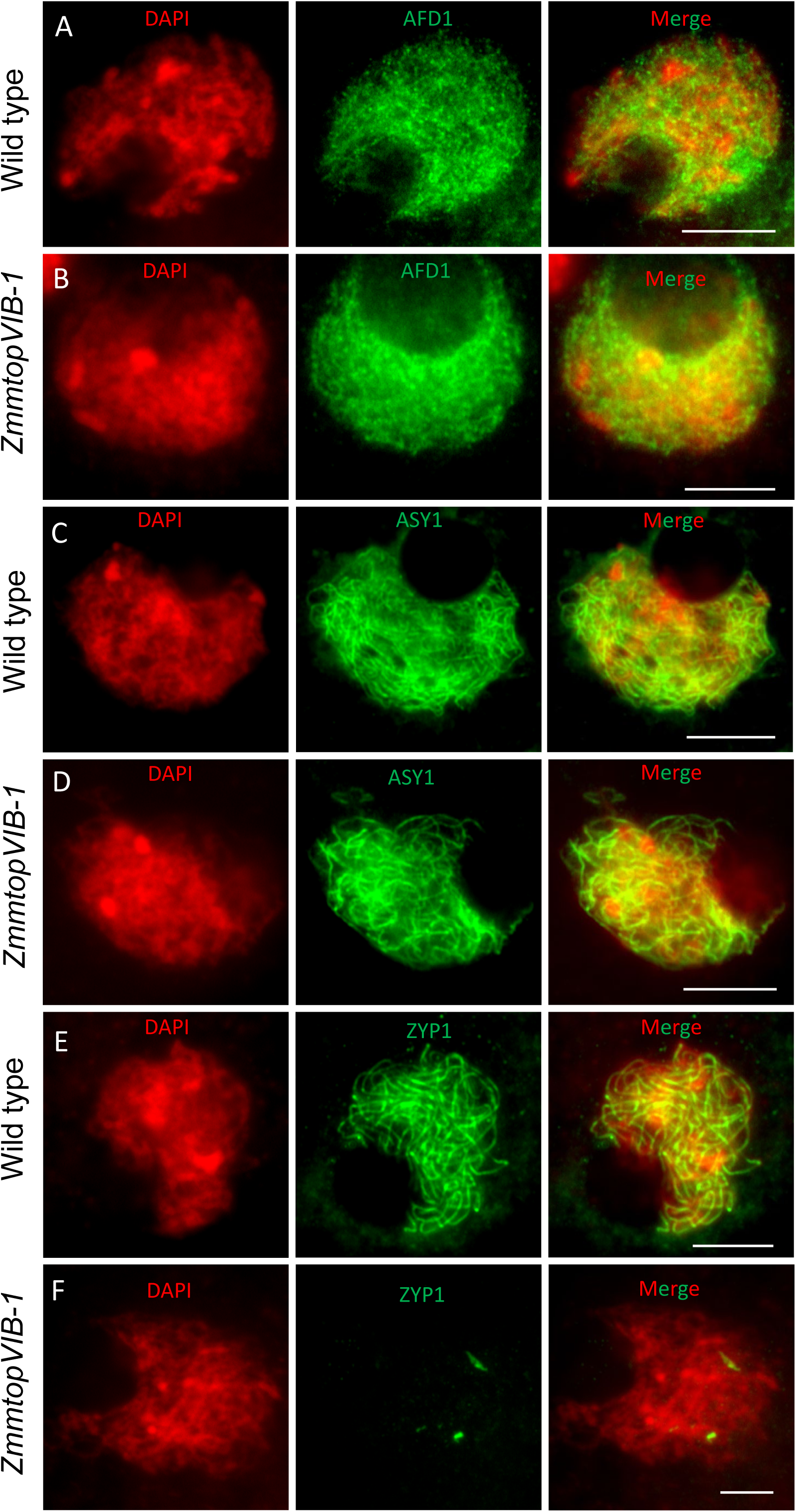
Immunofluorescence localization of AFD1, ASY1 and ZYP1 in wild type and *ZmmtopVIB-1* meiocytes. (A,C,E) AFD1 (A), ASY1 (C), and ZYP1 (E) in wild type meiocytes; (B,D,F) AFD1 (B), ASY1 (D), and ZYP1 (F) in *ZmmtopVIB-1* meiocytes. Scale bars = 10 µm.

Maize ASY1, a homolog of *Arabidopsis* ASY1 and rice PAIR2, is a basic element of AE and it is essential for SC assembly and HR (Armstrong et al., 2002; Golubovskaya et al., 2011). In wild type meiocytes, ASY1 loading manifested as linear signals along entire chromosomes in zygotene (Figure 6C, n=22). Again, ASY1 signals in *ZmmtopVIB-1* meiocytes were very similar to the pattern observed for wild type (Figure 6D, n=20), evidencing that *ZmMTOPVIB* is not required for AE installation.

Maize ZYP1, a homolog of *Arabidopsis* ZYP1 and rice ZEP1, is the central element (CE) of the SC and is assembled between AEs to regulate chromosome synapsis and CO formation (Higgins et al., 2005; Wang et al., 2010; Barakate et al., 2014). In wild type meiocytes, we observed elongated filaments of ZYP1 signals along the entire lengths of synapsed chromosomes at pachytene (Figure 6E, n=25). In contrast, only short branched or punctate ZYP1 signals were observed in *ZmmtopVIB-1* meiocytes at the same stage (Figure 6F, n=27), supporting that *ZmMTOPVIB* is crucial for ZYP1 loading. Taken together, these results indicate that *ZmMTOPVIB* is not required for AE installation, but it is indispensable for CE assembly during maize meiosis.

### *ZmMTOPVIB* is required for meiotic bipolar spindle assembly in maize

The recently identified function of *OsMTOPVIB* in regulating meiotic spindle assembly in rice prompted us to investigate if *ZmMTOPVIB* has the same role in maize meiosis. To do this, we performed immunostaining using α-tubulin antibody in wild type and *ZmmtopVIB-1* meiocytes. α-Tubulin heterodimerizes and polymerizes with β-tubulin to form microtubule walls (Blume et al., 2009; Gunning et al., 2015; Higgins et al., 2016). We found that microtubule filaments in wild type meiocytes gradually extended and attached to chromosomes during diakinesis (Figure 7A, n=13). At metaphase I, spindle fibers linked to the kinetochores and were arranged into two arrays, forming a canonical bipolar spindle structure (Figure 7B, n=25). At anaphase I, as the spindles extended, equal numbers of chromosomes were pulled by the bipolar spindles toward the opposite poles of the cell (Figure 7C, n=23). Finally, dyads formed, and the remaining spindles stayed at the equatorial plate (Figure 7D, n=18). In contrast, we consistently observed abnormal spindle structures in the *ZmmtopVIB-1* mutant at different stages of meiosis. In diakinesis, extension and aggregation of microtubule filaments was incomplete, with only one polar spindle forming so that the opposite cell pole lacked a polar spindle (Figure 7E and 7I, n=11). At metaphase I, spindle fibers of mutant meiocytes became entangled and distorted, lacking the typical bipolar spindle structure upon separation (Figure 7F and 7J, n=12). These distorted spindles dragged chromosomes randomly in multiple directions across the cell at anaphase I (Figure 7G and 7K, n=13), resulting in multinuclear dyads with abnormal equatorial plates in telophase I (Figure 7H and 7L, n=9).

**Figure 7.**
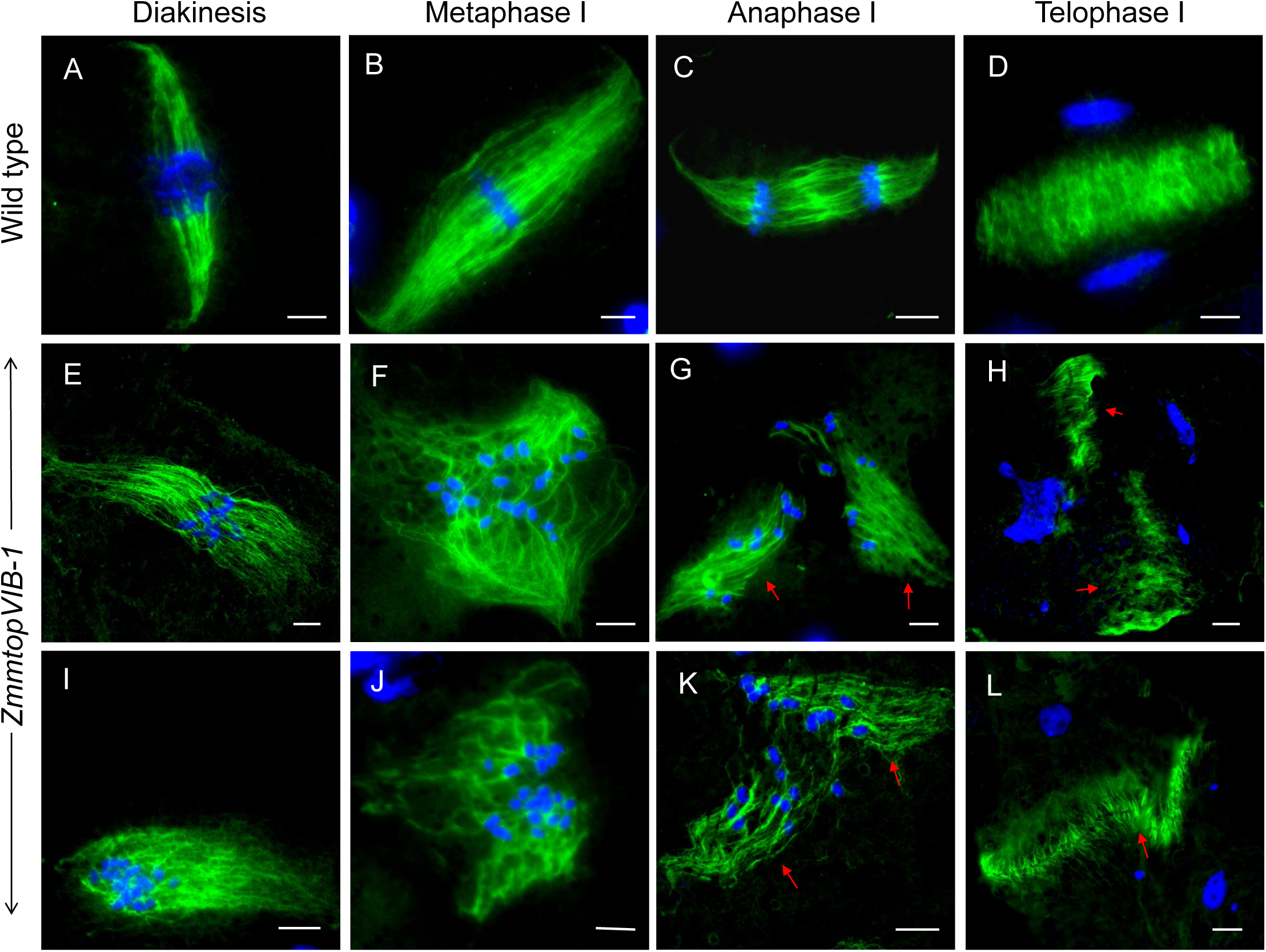
Meiotic spindle assembly process in wild type and *ZmmtopVIB-1.* (A-D) Spindle assembly in wild type. (E-L) Spindle assembly in *ZmmtopVIB-1.* (A,E,I) Diakinesis; (B,F,J) Metaphase I; (C,G,K) Anaphase I; (D,H,L) Telophase I. Chromosomes were marked with DAPI (blue), and microtubules were marked by α-tubulin antibody (green). The red arrows pointed out the abnormal spindles and equatorial plates. Scale bars = 10 μm.

To ascertain the outcomes of these abnormal spindle structures, we examined spore products in wild type and *ZmmtopVIB-1* mutant lines. In the wild type, we consistently observed symmetric dyads at telophase I (Figure 8A, n=55) and tetrads at telophase II (Figure 8E, n=52). In contrast, we frequently detected triads at telophase I (25.69%) (Figure 8C, n=28) and polyads at telophase II (63.24%) (Figure 8H, n=71) in *ZmmtopVIB-1*. Taken together, these results demonstrate that *ZmMTOPVIB* plays a critical role in proper bipolar spindle assembly during maize meiosis.

**Figure 8.**
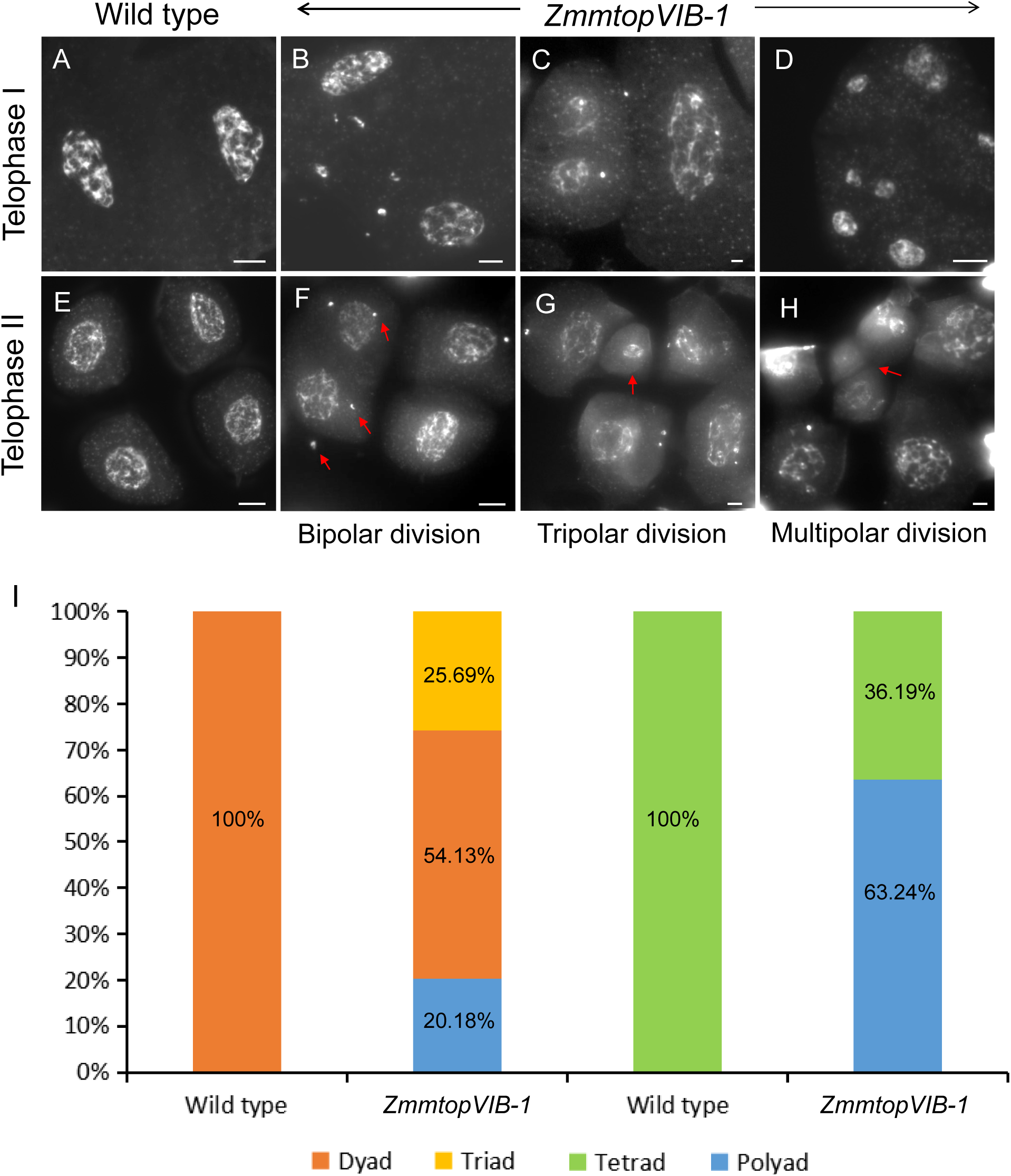
Meiotic products in wild type and *ZmmtopVIB-1*. (A,E) Meiotic cell division in wild type; (B,F) Bipolar division of *ZmmtopVIB-1* (B), producing tetrads (F); (C,G) Tripolar division of *ZmmtopVIB-1* (C), producing pentads/hexads (G); (D,H) Multipolar division of *ZmmtopVIB-1* (D), producing polyads (H); The red arrows pointed out the micronucleus and the abnormal additional spores. Scale bars=10μm. (I) Percentage of different meiotic products (dyad, triad, tetrad and polyad).

## Discussion

### Evolutionary conservation of *ZmMTOPVIB* function in meiotic DSB formation

We have shown that disruption of *ZmMTOPVIB* causes defects in the formation of meiotic DSBs and results in severe abnormalities of DSB-induced events, such as homologous synapsis, meiotic recombination and chromosome segregation. Although we detected few γH2AX signals in late zygotene stage, often seen to be associated with entangled chromosomes in mutant meiocytes, these sites are unlikely canonical meiotic DSBs. One possible scenario can be DNA damages resulted from chromosome knotting, which may be more frequent in a large genome with long chromosomes in the absence of recombination initiation. In support of this possibility, a similar observation was reported in maize *spo11-1* mutants (Ku et al., 2020). Nevertheless, our results demonstrate that *ZmMTOPVIB* is essential for normal meiotic DSB formation in maize, which are consistent with previous studies in other organisms, including mouse (Robert et al., 2016), *Arabidopsis* (Vrielynck et al., 2016) and rice (Fu et al., 2016; Xue et al., 2016), strengthening the notion that the role of TopoVIB in meiotic DSB formation is evolutionarily conserved among plants and mammals.

TopoVI is a heterotetramer comprised of two A subunits and two B subunits (Corbett and Berger, 2003; Gadelle et al., 2014). To generate DSBs on DNA, the A_2_ dimers catalyze the transesterification reaction for DNA cleavage (Bergerat et al., 1997), whereas the B_2_ subunits are responsible for the A_2_ conformation and DNA binding (Dutta and Inouye, 2000; Corbett et al., 2007; Graille et al., 2008). Although the basic set-up of the TopoVI complex seems conserved among organisms, the arrangement and operational mode of each component may vary. For instance, in mouse, the sole SPO11 gene produces two major SPO11 isoforms by alternative splicing, i.e. a short SPO11α and a longer SPO11β (Bellani et al., 2010). SPO11β interacts with TopoVIB to form a symmetrical heterotetramer for creating meiotic DSBs (Robert et al., 2016). In contrast, all land plants encode at least two SPO11 proteins, referred to as SPO11-1 and SPO11-2, both of which are required for DSB formation (Sprink and Hartung, 2014). Accordingly, the plant TopoVI complex is predicted to exist as an asymmetrical heterotetramer (Graille et al., 2008). Interestingly, *Arabidopsis* MTOPVIB not only acts as an essential component of the TopoVI complex, but also plays a critical role in mediating the formation of SPO11-1/SPO11-2 heterodimers, but not of SPO11-1 or SPO11-2 homodimers (Vrielynck et al., 2016; Jing et al., 2019). Hence, this function of *Arabidopsis* MTOPVIB as a linker to promote the assembly of SPO11 dimers may not necessarily be displayed by its counterpart in mouse. These results highlight that analogs of TopoVIB involved in meiotic DSB formation are evolutionarily conserved but subject to variation among different organisms.

### *ZmTOPVIB* is involved in meiotic bipolar spindle assembly

Proper morphogenesis and orientation of microtubule-based spindles are critical processes for ensuring the fidelity of chromosome segregation and cell division in eukaryotes (Gadde and Heald, 2004; Zhang and Dawe, 2011). In both monocot and dicot plants, construction of a meiotic bipolar spindle occurs by converting multipolar into bipolar spindle poles (Nannas and Dawe, 2016; Xue et al., 2019). However, the mechanism underlying meiotic bipolar spindle assembly remains unclear, with only a few contributory genes having been reported to date. *Arabidopsis ATK1* and *ATK5*, which encode kinesins, are motor proteins that typically advance along microtubules and presumably generate the force for microtubule movement, thereby affecting spindle organization (Chen et al., 2002; Ambrose and Cyr, 2007). *PRD2*, an essential gene for meiotic DSB formation in *Arabidopsis*, was originally named *Multipolar Spindle 1* (*MPS1*) based on the unequal bipolar or multipolar spindles present in *mps1* mutant meiocytes (Jiang et al., 2009). Intriguingly, OsMTOPVIB interacts with OsPRD2 in rice, indicating that these proteins may function in the same pathway, though the function of OsPRD2 has not yet been characterized (Xue et al., 2019). Since both *PRD2* and *OsMTOPVIB* are essential for DSB formation, it is conceivable that the processes of DSB formation and bipolar spindle assembly may be coordinated. However, this speculation was proven untrue upon discovery that bipolar spindles at metaphase I occur normally in three rice mutants exhibiting defects in DSB formation, i.e., the *pair2, Osspo11-1* and *crc1* mutants (Xue et al., 2019). Given these results, it is evident that meiotic spindle assembly is independent of DSB formation. Additionally, bipolar spindle assembly also seems to be uncoupled from homologous pairing since it was not perturbed in the *dsy1, dsy2*, and *afd1* mutants, three maize mutants defective in homologous pairing (Chan and Cande, 1998). Therefore, the molecular mechanism that integrates these HR-related genes in meiotic spindle assembly remains to be resolved.

As reported for the *OsmtopVIB* mutant, we observed a substantial proportion of triads at telophase I and polyads at telophase II in *ZmmtopVIB* meiocytes, demonstrating that maize *ZmMTOPVIB* is also required for meiotic bipolar spindle assembly. This finding evidences functional conservation of the *MTOPVIB* role in bipolar spindle assembly between rice and maize. However, we observed fewer abnormal triads or polyads in *ZmmtopVIB* than found in the *OsmtopVIB* mutant line. That discrepancy may be due to differential allelic effects on phenotypes. The two mutant alleles we considered in this study exhibited defective transcript splicing, hypothetically resulting in production of truncated protein. Although the complete abortion in DSB formation indicates that two alleles are at least severe hypomorphs, we cannot rule out the possibility that both *ZmmtopVIB* mutant alleles were not completely null for the specific role in bipolar spindle assembly, unlike for the previously studied *OsmtopVIB* mutant. Therefore, it would be worth investigating spindle defects in other *ZmmtopVIB* mutants in future studies.

Overall, our study demonstrates that *ZmMTOPVIB* is essential for meiotic DSB formation and that it plays a critical role in bipolar spindle assembly in maize. Our findings shed light on the evolutionary conservation of the dual functions of *MTOPVIB* in meiosis, though the mechanisms by which it plays a moonlighting role in bipolar spindle assembly await further investigation.

## Materials and Methods

### Plant material and genotyping

The UFMu-07260 mutant line (*ZmmtopVIB-1*) in the W22 inbred background was obtained from the UniformMu stock center of MaizeGDB (https://maizegdb.org/) (McCarty et al., 2013). Another mutant allele, EMS4-0742ae (*ZmmtopVIB-2*) in the B73 inbred background, was obtained from the Maize EMS induced Mutant Database (MEMD) (http://www.elabcaas.cn/memd/) (Lu et al., 2018). All plants were cultivated and fertilized under normal field condition during the growing season or in a growth chamber (16 h light at 28°C, 8 h dark at 22°C, 60-70% humidity). Maize genomic DNA was extracted using a method previously described (Li et al., 2013). Primers used for genotyping and sequencing of the two mutant alleles are listed in the Supplementary Table S1.

### Pollen viability

Pollen viability was assessed using the Alexander staining method (Alexander, 1969; Johnson-Brousseau and McCormick, 2004). Mature pollen grains were dissected out of anthers from wild type and *ZmmtopVIB* mutants during the pollination stage, and then stained with 10% Alexander solution. Images of stained pollen grains were taken using a Leica EZ4 HD stereo microscope equipped with Leica DM2000 LED illumination system (Leica, Solms, Germany).

### Real-time qPCR analysis

Total RNA was isolated from root, stem, leaf, developing embryo, endosperm, young tassel, and young ear using TRNzol-A^+^ Kit Reagent (TIANGEN, Beijing, China) according to the manufacturer’s instructions. Reverse transcription was performed using PrimeScriptTM II 1st strand cDNA Synthesis Kit (TaKaRa, Tokyo, Japan) with Oligo-T primer to obtain cDNA. Quantitative PCR was conducted with a 7500 Fast Real-Time PCR System (Applied Biosystems, Foster City, CA, USA) using SYBR Green Master Mix (TaKaRa). All reactions were performed with three biological replicates and technical repeats. Gene-specific primers and reference gene (*Ubiquitin*) primers for internal control are listed in the Supplementary Table S1.

### Meiotic chromosome preparation and DAPI staining

Young tassels were fixed in Carnoy’s solution (ethanol: glacial acetic acid; 3:1) for 1 day at room temperature, and then stored in 70% ethanol at 4 °C. Anthers were dissected in 45% acetic acid solution. Meiocytes were squeezed from anthers and squashed onto slides using coverslips. Slides were frozen in liquid nitrogen and the coverslips were removed immediately. After serial dehydration in 70%, 90% and 100% ethanol, the air-dried slides were stained and mounted with 4, 6-diamidinophenylindole (DAPI) in Vectashield anti-fade medium (Vector Laboratories, CA, USA).

### FISH and chromosome painting

Chromosome spreads were prepared by the method described previously (Wang et al., 2006). Three repetitive DNA probes were used, including the pTa794 clone containing 5S rDNA repeats (Li and Arumuganathan, 2001), the pAtT4 clone containing telomere-specific repeats (Richards and Ausubel, 1988), and cy5-conjugated 180-bp knob oligonucleotides. The rDNA and telomere probes were labeled by nick translation kit (Roche, Basel, Switzerland). The chromosome 3 painting probe was labeled with ATTO-550 as previously described (Albert et al., 2019). Slides were counterstained using DAPI in anti-fade mounting medium (Vector Laboratories). Chromosome images were captured under a Ci-S-FL fluorescence microscope (Nikon) equipped with a DS-Qi2 microscopy camera (Nikon, Tokyo, Japan) or using a Delta Vision ELITE system (GE) equipped with an Olympus IX71 microscope.

### Immunofluorescence assay

Immunofluorescence was performed as described previously (Pawlowski et al., 2003) with minor modifications. After being dissected and permeabilized in 1×Buffer A solution with 4% (w/v) paraformaldehyde for 30 min at room temperature, fresh young anthers were washed twice in 1×Buffer A at room temperature, and then stored in 1×Buffer A at 4 °C. Meiocytes were squeezed from anthers and squashed onto slides. After frozen in liquid nitrogen, coverslips were removed immediately. The meiocytes were incubated in blocking buffer diluted with primary antibodies for 1 hour in a 37 °C humidity chamber, then washed three times in 1× PBS. Goat anti-rabbit antibodies conjugated with Fluor 555 diluted in blocking buffer were added to the slides. After incubation at 37°C for 1 hour, the slides were washed three times in 1× PBS. Finally, cells were counterstained with DAPI in anti-fade mounting medium (Vector Laboratories). The antibodies against ASY1, ZYP1, and γH2AX were prepared in our laboratory as described previously (Jing et al., 2019). Antibodies against AFD1, RAD51, and DMC1 were gifts from collaborators. All primary and secondary antibodies were diluted at 1:100. Images of meiocytes were observed and captured using a Ci-S-FL microscope (Nikon) equipped with a DS-Qi2 microscopy camera (Nikon, Tokyo, Japan). 2D projected images were generated using NIS-Elements software. Further image processing was conducted using ImageJ software (https://imagej.nih.gov/ij/index.html).

## Acknowledgments

We thank all members of our laboratories for helpful discussion and assistance during this research. We greatly appreciate Changbin Chen (University of Minnesota), Wojciech Pawlowski (Cornell University), and Huabang Chen (Chinese Academy of Science, China) for gifting us AFD1, RAD51, and DMC1 antibodies, respectively. We are also grateful to James Birchler (University of Missouri) for generous gifts of chromosome-specific painting probes. We thank the cell biology core laboratory in IPMB, Academia Sinica for microscope imaging technical support.

